# Vertically transmitted rhabdoviruses are found across three insect families and have dynamic interactions with their hosts

**DOI:** 10.1101/084558

**Authors:** Ben Longdon, Jonathan P Day, Nora Schulz, Philip T Leftwich, Maaike A de Jong, Casper J Breuker, Melanie Gibbs, Darren J Obbard, Lena Wilfert, Sophia CL Smith, John E McGonigle, Thomas M Houslay, Lucy I Wright, Luca Livraghi, Luke C Evans, Lucy A Friend, Tracey Chapman, John Vontas, Natasa Kambouraki, Francis M Jiggins

## Abstract

A small number of free-living viruses have been found to be obligately vertically transmitted, but it remains uncertain how widespread vertically transmitted viruses are and how quickly they can spread through host populations. Recent metagenomic studies have found several insects to be infected with sigma viruses (*Rhabdoviridae*). Here, we report that sigma viruses that infect Mediterranean fruit flies (*Ceratitis capitata*), *Drosophila immigrans*, and speckled wood butterflies (*Pararge aegeria*) are all vertically transmitted. We find patterns of vertical transmission that are consistent with those seen in *Drosophila* sigma viruses, with high rates of maternal transmission, and lower rates of paternal transmission. This mode of transmission allows them to spread rapidly in populations, and using viral sequence data we found the viruses in *D. immigrans* and *C. capitata* had both recently swept through host populations. The viruses were common in nature, with mean prevalences of 12% in *C. capitata,* 38% in *D. immigrans* and 74% in *P. aegeria*. We conclude that vertically transmitted rhabdoviruses may be widespread in insects, and that these viruses can have dynamic interactions with their hosts.

## Introduction

Insects are host to a range of vertically transmitted parasites [1, 2]. Vertical transmission is normally associated with maternally transmitted bacterial endosymbionts, and with transposable elements that can proliferate within the host genome and spread through populations [3, 4]. Many free-living viruses are also capable of vertical transmission without integrating into the host genome [5]. In most cases this is combined with horizontal transmission [6], but in a few cases the virus has become obligately vertically transmitted [2, 7].

A well characterised obligately vertically transmitted virus that does not integrate into its host’s genome is the sigma virus of *Drosophila melanogaster* (DMelSV) [8]. This is a negative sense RNA virus in the family *Rhabdoviridae* that is found in the cytoplasm of cells [9]. Typically, vertically-transmitted cytoplasmic pathogens can only be transmitted maternally, and this alone cannot allow them to increase in prevalence [10]. However, DMelSV is transmitted by both infected males and females, which allows it to spread through host populations, even if it carries a cost to the host [2, 9, 11–13]. This is analogous to the spread of transposable elements through a host population, where the element can spread at rates greater than a Mendelian locus, giving it a natural transmission advantage. Two more vertically transmitted sigma viruses have been characterised in different species of *Drosophila* (DAffSV and DObsSV from *Drosophila affinis* and *Drosophila obscura* respectively). Like DMelSV, these are biparentally transmitted through both the eggs and sperm of their hosts [14, 15]. Females transmit the virus at a high rate (typically close to 100%), whereas males transmit the virus at a lower rate, probably because sperm transmit a lower amount of virus to the developing embryo [9, 15].

A consequence of biparental transmission is that, like transposable elements, sigma viruses can rapidly spread through host populations [3,4, 16]. For example, a genotype of DMelSV that was able to overcome a host resistance gene called *ref(2)P* swept across Europe in the 1980-1990s [17–20]. Similarly, DObsSV has swept through populations of *D. obscura* in the last decade [15].

It is unknown whether obligately vertically transmitted viruses are common in nature or are just a quirk of a few species of *Drosophila.* Recent metagenomic sequencing has found rhabdoviruses associated with a wide diversity of insects and other arthropods, including numerous viruses closely related to the three vertically transmitted sigma viruses described in *Drosophila* [21, 22]. This raises the prospect that vertically transmitted rhabdoviruses could be common insect pathogens. Here, we examine three recently identified viruses that fall into the sigma virus clade and infect Mediterranean fruit flies (medflies; *Ceratitis capitata*, Diptera, Tepthritidae), *Drosophila immigrans* (Diptera, Drosophilidae, sub-genus Drosophila) and speckled wood butterflies (*Pararge aegeria,* Lepidoptera, Nymphalidae). We go on to test whether these viruses are vertically transmitted and investigate whether they show evidence of the rapid dynamics seen in other sigma viruses.

## Methods

### Transmission

We determined the patterns of transmission of *Ceratitis capitata sigmavirus* (CCapSV), *Drosophila immigrans sigmavirus* (DImmSV) and *Pararge aegeria rhabdovirus* (PAegRV), which all fall into the sigma virus clade [21]. We carried out crosses between infected and uninfected males and females, and measured the rates of transmission to their offspring.

Infected *C. capitata* were collected from the Cepa Petapa lab stock and uninfected flies were from the TOLIMAN lab stock. Virgin females and males were crossed and their offspring collected. Only 29% of flies from the Cepa Petapa stock were infected when we carried out the crosses (see below). In total we tested 10 crosses between infected females and uninfected males, 8 crosses with uninfected females and infected males, and 7 crosses where neither sex was infected. We tested both parents and a mean of 6 offspring for each cross (range 4-8, total of 197 offspring) for infection using RT-PCR (see table S1).

Infected *D. immigrans* were collected from a DImmSV infected isofemale line (EGL 154) and uninfected flies were collected from a stock established from four isofemale lines (all lines originated from Cambridge UK). Virgin females and males were crossed and their offspring collected. In total we tested 20 crosses between infected females and uninfected males, 18 crosses with uninfected females and infected males, and 8 crosses where neither sex was infected. We tested both parents and a mean of 4 offspring for each cross (range= 2-4, total of 178 offspring) for infection using RT-PCR (see table S1).

To measure transmission of PAegSV in speckled wood butterflies, we examined crosses between the offspring of wild caught *P. aegeria* females that were an unknown mix of infected and uninfected individuals. The wild caught females were collected in Corsica and Sardinia in May 2014 (see below). Virgin females and males were crossed, and their offspring collected. We tested the infection status of the parents used for the crosses *post hoc* using RT-PCR; in total there were 10 crosses between infected females and uninfected males, 8 crosses with uninfected females and infected males and 1 cross where neither sex was infected (data not shown from 28 crosses where both parents were infected). We tested a mean of 4 offspring for each cross (range 1-8, total of 171 offspring) for virus infection using RT-PCR (see table S1).

### Collections to examine virus prevalence in wild populations

*C. capitata* were collected (as eclosing flies) from fallen argan fruit in Arzou, Ait Melloul, Morocco in July 2014 (latitude, longitude: 30.350, −9.473) and from peaches and figs from Timpaki, Crete from July-September 2015 (35.102, 24.756).

*D. immigrans* were collected from August-October 2012 from the following locations: Kent (51.099, 0.164 and 51.096, 0.173); Edinburgh (55.928, −3.169 and 55.925, −3.192); Falmouth (50.158, −5.076 and 50.170, −5.107); Coventry (52.386, −1.482; 2.410, −1.468; 52.386, −1.483 and 52.408, −1.582); Cambridge (52.221, 0.042); Derbyshire (52.978, −1.439 and 52.903, −1.374); Les Gorges du Chambon, France (45.662, 0.555) and Porto, Portugal (41.050, −8.645).

*P. aegeria* were collected from several UK locations in August and September 2014: South Cambridgeshire (52.116, 0.252); Yorkshire (53.657, −1.471); Oxfordshire (51.833, −1.026); North Cambridgeshire (52.395, −0.237); Dorset (50.999, −2.257) and Somerset (51.363, −2.525). We also sequenced one infected *P. aegeria* individual from each of the families used for the crosses described above. These individuals were collected in May 2014 from Corsica (41.752, 9.191; 41.759, 9.184; 41.810, 9.246; 41.377, 9.179; 41.407, 9.171; 41.862, 9.379; 42.443, 9.011 and 42.516, 9.174) and Sardinia (39.964, 9.139; 39.945, 9.199; 41.233, 9.408; 40.911, 9.095 and 40.037, 9.256).

### Virus detection and sequencing

Individual insects were homogenised in Trizol reagent (Invitrogen) and RNA extracted by chloroform phase separation, followed by reverse transcription with random–hexamers using GoScript reverse transcriptase (Promega). For each sample we carried out PCRs to amplify partial nucleocapsid (N) and RNA Dependant RNA Polymerase (L) gene sequences from the respective viral genomes (see table S1), as well as a control gene from the insect genome (*COI* or *RpL32*) to confirm the extraction was successful. For CCapSV and PAegRV we sequenced all infected samples (19 and 130 respectively), for DImmSV we sequenced a subset of 87 samples from across all populations. PCR products were treated with Antarctic Phosphatase and Exonuclease I (New England Biolabs) and directly sequenced using BigDye on a Sanger ABI capillary sequencer (Source Bioscience, Cambridge, UK). Data were trimmed in Sequencher (v4.5) and aligned using ClustalW in Bioedit software. All polymorphic sites were examined by eye. Recombination is typically absent or rare in negative sense RNA viruses [23]; we were unable to detect any evidence of recombination in our data using GARD [24] with a general time reversible model and gamma-distributed rate variation with four categories. Median joining phylogenetic networks were produced using PopArt (v1.7 http://popart.otago.ac.nz.). Population genetic analysis was carried out in DNAsp (v5.10.01). *P* values for estimates of Tajima’s *D* were estimated using DNAsp; across all sites *P* values were calculated using coalescent simulations assuming no recombination, for synonymous sites they were estimated using a beta distribution. Maps used for the figures were from QGIS (v2.14) [25].

### ADAR edits

We observed some of the DImmSV sequences had a cluster of mutations in the N gene consistent with those caused by adenosine deaminases that act on RNAs (ADARs). ADARs target double stranded RNA and convert adenosine (A) to inosine (I), and display a 5’ neighbour preference (A=U>C>G) [26, 27]. During viral genome replication I’s are paired with guanosine (G), so editing events appear as changes from A to G when sequenced. Sigma viruses have been found to show mutations characteristic of ADAR editing, with single editing events causing clusters of mutations [28, 29].

As ADAR-induced hyper-mutations will be a source of non-independent mutation we aimed to exclude such mutations to prevent them confounding our analyses. Compared to a 50% majority-rule consensus sequence of our DImmSV sequences, we identified 17 sites with A to G mutations at ADAR preferred sites across the negative sense genome and its positive sense replication intermediate. This is a significant over-representation when compared to ADAR non-preferred sites (Fisher’s exact test *P*=0.037). We therefore excluded all ADAR preferred sites from our dataset and carried out our population genetics analysis on this data. For CCapSV and PAegRV sequences we did not detect an overrepresentation of A to G mutations at ADAR preferred sites, suggesting these viruses did not contain ADAR hyper-mutations. The R [30] script used to identify ADAR edits and exclude preferential ADAR editing sites is available in a data repository (https://dx.doi.org/10.6084/m9.figshare.3438557.v1).

### Reconstructing viral population history

To reconstruct how long ago these viruses shared a common ancestor and how their population size has changed over time we used a Bayesian phylogenetic inference package (BEAST v1.8.0) [31]. The evolutionary rate of viruses was assumed to be the same as in DMelSV [32]. This is similar to other related rhabdoviruses [33, 34] and evolutionary rates of DMelSV do not differ significantly between the lab and field [35]. To account for uncertainty in this evolutionary rate estimate, we approximated its distribution with a normal distribution (mean=9.9×10-5 substitutions/site/year, standard deviation=3.6×10^-5^, substitutions/site/year), and this distribution was used as a fully-informative prior to infer dates of the most recent common ancestor of each virus. The model assumed a strict molecular clock model and an HKY85 substitution model [36]. Sites were partitioned into two categories by codon position (1+2, 3), and separate evolutionary rates were estimated for each category. Such codon partition models have been shown to perform as well as more complex non-codon partitioned models but with fewer parameters [37]. We tested whether a strict clock rate can be excluded by running models with a lognormal relaxed clock; for all three viruses we found the posterior estimate of the coefficient of variation statistic abuts the zero boundary, and so a strict clock cannot be excluded [38].

We reconstructed the phylogeny with an exponentially expanding population (parameterised in terms of growth rate) or a constant population size model. The population doubling time was calculated from the growth rate as ln(2)/growth rate. We excluded a constant population size if the 95% confidence intervals (CIs) for estimates of growth rate did not cross zero. We verified we were using the most suitable demographic model for each virus using the path sampling maximum likelihood estimator implemented in BEAST (see supplementary materials) [39]. We note that estimates of the root age were similar across models (Table S2).

Each model was run for 1 billion MCMC steps with sampling every one hundred thousand generations for each model, and a 10% burnin was used for all parameter estimates, selected after examining trace files by eye. Posterior distributions and model convergence was examined using Tracer (v1.6) [40] to ensure an adequate number of independent samples. The 95% CI was taken as the region with the 95% highest posterior density.

## Results

### Sigma viruses in speckled wood butterflies, medflies and *Drosophila immigrans* are vertically transmitted

The sigma virus of *D. melanogaster* is a vertically transmitted through both eggs and sperm. In a recent metagenomics analysis we identified sigma viruses in a range of other insect species [21]. Here, we report that three of these viruses are also vertically transmitted, with large sex-differences in transmission rates. In all three species the infected females transmitted the virus to the majority of their offspring (Figure 1; mean proportion offspring infected: CCapSV=0.82, DImmSV=1, PAegRV=0.88). Paternal transmission rates were much lower (Figure 1). Infected male *C. capitata* did not transmit CCapSV to any of their offspring. Infected male *D. immigrans* transmitted DImmSV to their offspring, but at lower rate (0.51) than through females (Wilcoxon exact rank test: *W*=20, *P*<0.001). Similarly, infected male *P. aegeria* also transmitted PAegRV to their offspring, but at lower rate (0.51) than maternal transmission (Wilcoxon rank sum test *W*=17, *P*=0.026). There was no difference in the proportion of infected sons and daughters for all three viruses (Wilcoxon exact rank test: CCapSV *W*=316, *P*=1; DImmSV *W*=1106, *P*=0.538, PAegRV *W*=782, *P*=0.853).

**Figure 1.**
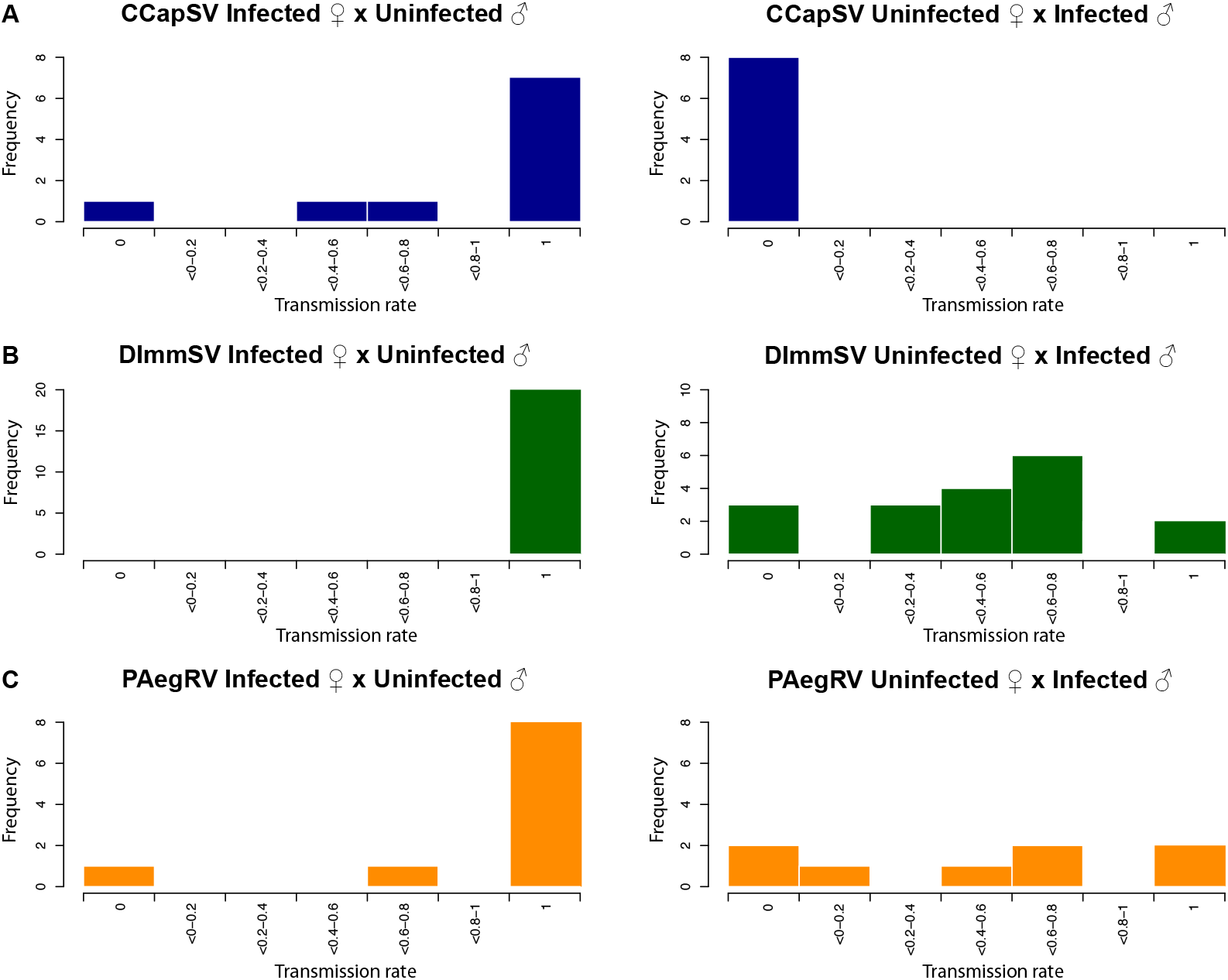
Vertical transmission rates of three sigma viruses from infected females (left) and males (right). A= CCapSV, B= DImmSV, C= PAegRV. The far left and far right bins are individuals with zero or 100% transmission respectively. Results from control crosses where both parents were uninfected are not shown.

For CCapSV and DImmSV horizontal or sexual transmission appears to be rare or absent. We did not detect virus in any of the offspring from crosses where both parents were uninfected, or in uninfected individuals that mated with infected individuals during the crosses. As the infection status of parents in crosses to measure transmission of PAegRV was only established *post hoc*, we were unable to test for evidence of horizontal or sexual transmission between parents.

### Sigma viruses are common in natural populations

We tested 243 *C. capitata*, 527 *D. immigrans* and 137 *P. aegeria* from the wild for presence of their respective viruses using RT-PCR. We found the mean viral prevalence across populations was 12% for CCapSV, 38% for DImmSV and 74% for PAegRV. There were significant differences in the prevalence between populations (tables S4-S6) for CCapSV (Figure 2; Chi-Sq test, df = 1, *χ*^2^= 13.08, *P* =<0.001) and DImmSV (Figure 2; Chi–Sq test, df=7, *χ*^2^=40.648, *P*<0.001), but not for PAegSV (Figure 2; Chi-Sq test, df = 72, *χ*^2^= 81.333, *P*=0.211).

**Figure 2.**
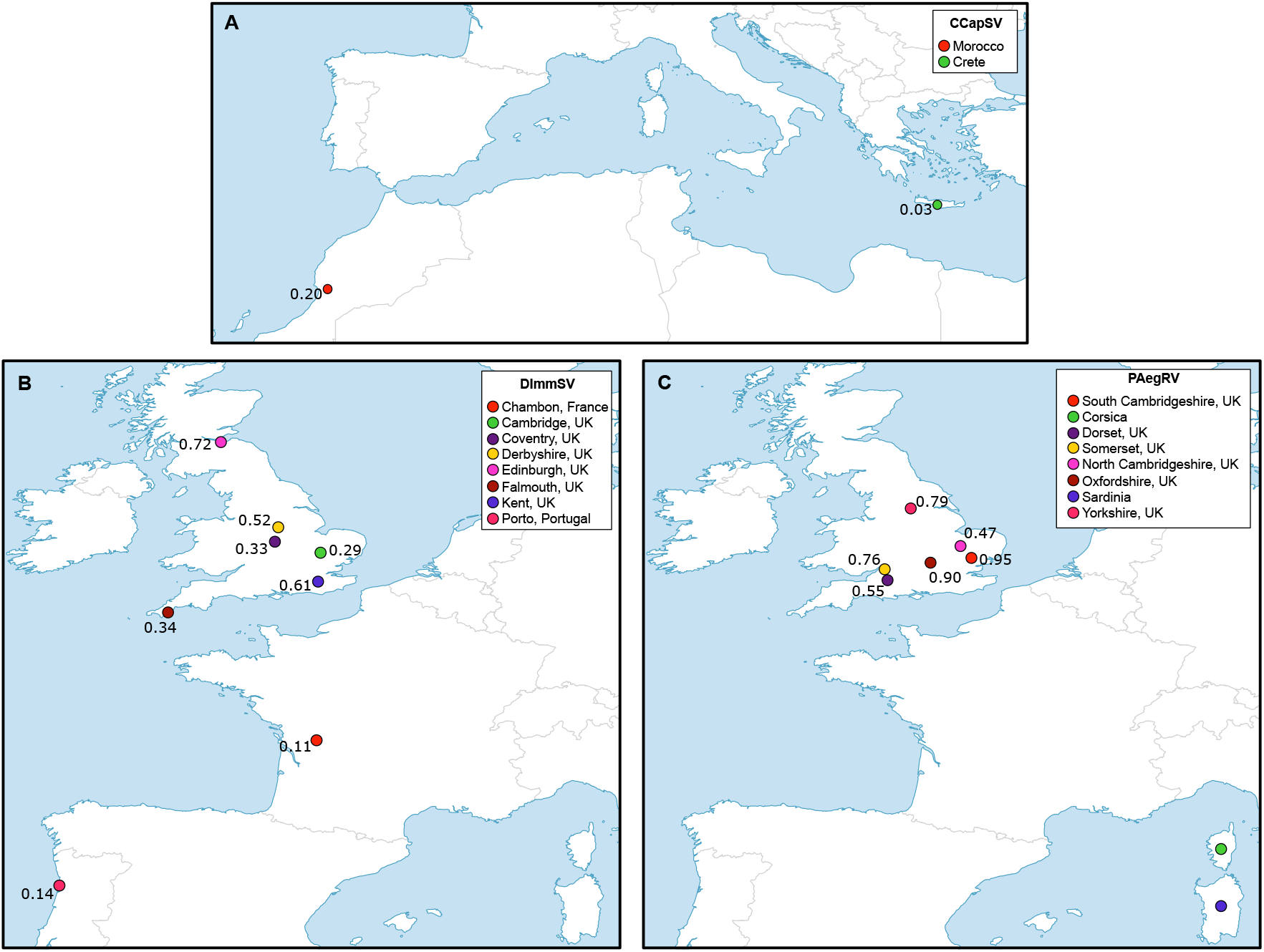
Viral prevalence at different locations. A= CapSV; B= DImmSV and C= PAegRV. Prevalence data was not available for PAegRV collected in Corsica and Sardinia.

### Sigma viruses can rapidly spread through populations

To infer the past dynamics of these viruses, we sequenced part of the N and L genes from multiple viral isolates. We used these sequence data to produce a median joining phylogenetic network for each of the three viruses (Figure 3), and examined the population genetics of these virus populations.

DImmSV had the lowest genetic diversity of the three viruses, and appears to have very recently swept through host populations. We found only 40 polymorphic sites out of 929 sites examined (16 in N and 24 in L gene) over 87 viral sequences. The average number of pairwise differences per site (π) was 0.20% across all sites. This low genetic diversity appears to have been caused by the virus recently sweeping through host populations, with the DImmSV sequences forming a star shaped network as is expected following a recent sweep (Figure 3). Due to the low levels of population structure and small sample sizes from individual populations (see below), we combined sequences from across populations to investigate the past demography of the virus. Overall there was a large excess of rare variants compared to that expected under the neutral model, which is indicative of an expanding population or selective sweep (Tajima’s *D*= −2.45, *P*<0.001). Out of 40 segregating sites, 27 are singletons. This result held even if only synonymous sites were analysed (Tajima’s *D*=-2.27 *P*<0.01), indicating that it is not likely to be caused by slightly deleterious amino acid polymorphisms being kept at low frequency by purifying selection. Furthermore, these results are unlikely to be confounded by population structure, as when we analysed only the samples from Derbyshire (the population with the largest sample size, n=41) we found Tajima’s *D* was significant for all sites (*D*= −1.68 *P*=0.026). Tajima’s *D* was negative but not significant for synonymous sites (synonymous: *D*=-1.11 *P*>0.1), probably because there were only six synonymous polymorphisms in this population.

To reconstruct past changes in the effective size of the DImmSV population we used a Bayesian approach based on the coalescent process in the BEAST software [31, 41]. The posterior distribution for the estimated growth rate did not overlap zero (95% CI = 0.12, 0.99) suggesting the population had expanded, and the exponential growth model was also preferred in the path sampling analysis (see table S3). Assuming the evolutionary rate is the same as the related *D. melanogaster* virus DMelSV, we estimated the viral population size has doubled every 1.5 years (95% CI= 0.7-5.7), with the viruses in our sample sharing a common ancestor 16 years ago (95% CI= 5-31 years).

CCapSV had higher genetic diversity, likely reflecting a somewhat older infection than DImmSV. Combining sequences across populations, the genetic diversity was approximately five times greater than DImmSV (? = 0.99% across all sites) and we found 44 segregating sites over 1278 sites (21 in N gene and 23 in L gene) in 19 viral sequences. As CCapSV showed high levels of genetic population structure (see results below), we restricted our analyses of demography to viruses from Morocco (the population with the greatest number of samples, n=13). For these viruses we found a significant excess of rare variants at all sites (Tajima’s *D*= −1.67 *P*=0.035) and for synonymous sites (Tajima’s *D*=-1.81 *P*<0.05). This suggests the Moroccan population of CCapSV has been expanding or undergone a recent selective sweep. The coalescent analysis supported the hypothesis that the CCapSV population had expanded (95% CI of the exponential growth parameter = 0.031, 0.343) and the exponential growth model was preferred in the path sampling analysis (see table S3). We estimated the effective population size has doubled every 3.9 years (95% CI= 2.0-22.7 years), with the Moroccan viruses sharing a common ancestor 45 years ago (95% CI=16-85 years) suggesting this is a more ancient expansion, with a greater number of mutations accumulating since the lineages separated. Combining the samples from the two populations, the most recent common ancestor of all CCapSV isolates was estimated to be 51 years ago (95% CI=18-96 years) with an exponential growth model, assuming the evolutionary rate is the same as DMelSV.

PAegRV was inferred to be the oldest of the three infections. The genetic diversity of this virus was over 11 times that of DImmSV (?: 2.22% across all sites), and we found 204 segregating sites over 1281 sites (83 in N gene 121 in L gene) in 130 viral sequences. As PAegRV showed high levels of genetic population structure (see results below), we restricted our analyses of demography to viruses from Hildersham (the population with the largest sample size, n=36). There was no evidence of a population expansion in viruses from Hildersham, as there was not an excess of rare variants for all sites (Tajima’s *D*= −0.58, *P*=0.322) or synonymous sites (Tajima’s *D*= −0.65, *P*>0.1). Furthermore, we could reject a model of exponential growth in BEAST, as the 95% CI of the growth rate overlapped zero (95% CI= −0.008, 0.028), and the constant population size model was preferred in the path sampling analysis (see table S3). The BEAST analysis supported the conclusion that this was an older infection, with the common ancestor of all of our viral isolates existing 309 years ago (95% CI=105-588 years), assuming the evolutionary rate is the same as DMelSV.

**Figure 3:**
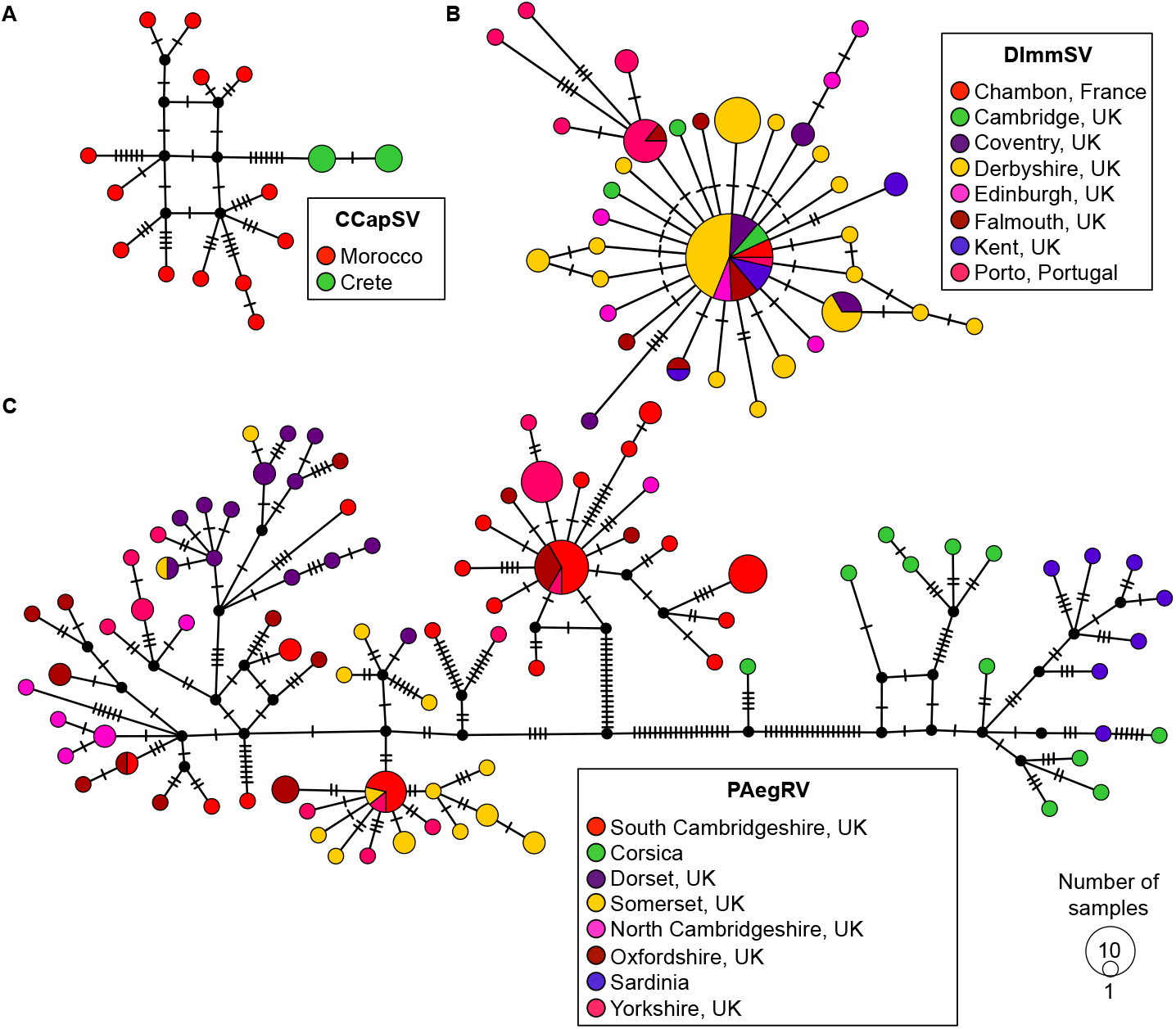
Median joining phylogenetic network of sequences from the three viruses. The colours represent the different locations samples were collected from, the size of the node represents the number of samples with that sequence and the dashes on branches show the number of mutations between nodes. A= 19 CCapSV sequences, B= 87 DImmSV sequences, C= 130 PAegRV sequences. Phylogenetic trees of each of the viruses are also available (https://dx.doi.org/10.6084/m9.figshare.3437723.v1).

### Sigma viruses have genetically structured populations

The virus populations were all geographically genetically structured, with the oldest infections (see above) showing the greatest levels of structure. Using partial sequences of the N and L genes, we quantified genetic structure by calculating an analogue of *FST* (*KST*) that measures the proportion of the genetic variation contained in subpopulations relative to the population as a whole [42]. The eight PAegSV populations showed high levels of genetic differentiation (Figure 3), with *KST* values of 0.48 (permutation test, *P*<0.001). Even when the divergent Corsican and Sardinian populations were excluded we still found significant genetic differentiation (*KST* =0.24, permutation test, *P*<0.001). The two CCapSV populations fell into monophyletic clades, and this was reflected in intermediate levels of genetic differentiation (Figure 3) with *KST* = 0.37 (permutation test, *P*<0.001). Finally, DImmSV showed lower levels of genetic differentiation with a *KST* value of 0.15 (permutation test, *P*<0.001).

## Discussion

Free-living viruses that are obligately vertically transmitted have only been reported from a small number of insect species. However, recent metagenomic studies have found that sigma viruses related to a vertically transmitted pathogen of *Drosophila melanogaster* are widespread in insects. Here we have sampled three of these viruses from different insect families (Lepidoptera, and Diperta in the Tephritidae and Drosophilidae), and found that they are all vertically transmitted. Therefore, most sigma viruses are likely vertically transmitted, and may represent a major group of insect pathogens.

The patterns of vertical transmission that we observed for three sigma viruses from *C. capitata*, *D. immigrans* and *P. aegeria* are consistent with the mode of vertical transmission seen in other sigma viruses [9, 15]. In two of the viruses we studied — PAegRV and DImmSV — we found transmission through both eggs and sperm, but higher transmission rates through eggs. However, we only observed maternal transmission of CCapSV. Paternal transmission may also occur in CCapSV but we simply failed to detect it in this experiment, as in other sigma viruses some infected males are unable to transmit the virus if the line is “unstabilised” [9, 15]. This is commonly seen in insect lines with low infection rates (only 29% of flies carried CCapSV in our infected line) and appears to be due to males that are infected from their father receiving only a low dose of virus that fails to infect the male germ-line [9, 15].

Our population genetic data shows this mode of transmission can allow rapid sweeps of these viruses through host populations. Many cytoplasmic bacteria are also vertically transmitted in insects, and these are almost exclusively only transmitted by infected females. This has led these endosymbionts to evolve different strategies to ensure their persistence, such as distorting the host sex ratio and causing cytoplasmic incompatibility [10]. Biparental transmission is an alternate strategy that can allow sigma viruses to rapidly invade host populations [32, 43]. Here, we have shown DImmSV and CCapSV sequences both show evidence of recent sweeps (~15 and 50 years ago respectively). Of the two sigma viruses studied previously (DMelSV from *D. melanogaster* and DObsSV in *D. obscura*) both showed evidence of a recent sweep [15, 32, 35]. The spread of DImmSV and CCapSV has occurred in the last few decades, with the viral populations doubling in just a few years. Therefore, sigma viruses seem to have very dynamic associations with host populations. This reflects the pattern seen in other vertically transmitted parasites. For example P-elements invaded populations of *D. melanogaster* worldwide in the twentieth century and are currently spreading through *D. simulans* populations [3, 16]. Likewise, vertically transmitted bacterial endosymbionts have frequently been found to have recently swept through new species or populations [44–47].

Why do sigma viruses have such dynamic interactions with their hosts? In the case of DMelSV this was thought to be a selective sweep of viral genotypes able to overcome a host resistance gene called *ref(2)P* [19, 20, 48]. DMelSV genotypes that could overcome the *ref(2)P* resistance allele rapidly increased in frequency between the early 1980s and early 1990s in French and German populations [18, 49]. In a recombining population a selective sweep would only reduce diversity surrounding the site under selection, but as these viruses do not recombine the entire genome is affected by a selective sweep. For DImmSV, CCapSV and DObsSV we are therefore unable to distinguish between a long–term host-virus association overlain by a recent selective sweep of an advantageous mutation, or the recent acquisition and spread of novel viruses through previously uninfected host populations. To separate these hypotheses, we would need to discover either closely related viruses in other species or populations, or the remnants of more diverse viral populations that existed prior to a selective sweep.

## Conclusions

Our results suggest that vertically transmitted rhabdoviruses may be widespread in a broad range of insect taxa. It remains to be seen whether this mode of vertical transmission is a unique trait of sigma-like rhabdoviruses, or whether this is the case for the numerous rhabdoviruses from other clades that infect insects [21, 22]. Sigma viruses commonly have dynamic interactions with their hosts, with vertical transmission though both eggs and sperm enabling them to rapidly spread through host populations.

## Author contributions

Conceived and designed study: BL. Carried out field collections: BL, SCLS, DJO, JEM, LIW, MDJ, CJB, MG, TMH, PL, JV, NK. Carried out crosses to measure transmission: PL, TC, LAF, CJB, MG, LL, LCE, SCLS, JPD, BL. Carried out molecular work: NS, JPD, SCLS, BL. Provided resources/samples for project: all authors. Analysed data: BL and FMJ. Wrote manuscript: BL and FMJ with comments from all other authors.

## Data availability

Sequences are deposited in Genbank under the following accessions: KX352837–KX353310

Additional data are available on Figshare:

Sequence alignments (https://dx.doi.org/10.6084/m9.figshare.3409420.v1)

ADAR R script (https://dx.doi.org/10.6084/m9.figshare.3438557.v1)

Phylogenetic trees (https://dx.doi.org/10.6084/m9.figshare.3437723.v1)

Transmission data (https://dx.doi.org/10.6084/m9.figshare.4133469.v1)

## Acknowledgements

Thanks to Roger Vila, Leonardo Dapporto, Vlad Dinca and Raluca Voda who collected the Sardinian and Corsican *P. aegeria* to set-up the lab stocks used for crosses.

